# Diagnostic Whole Exome Sequencing in Patients with Short Stature

**DOI:** 10.1101/414987

**Authors:** Huijuan Zhu, Ziying Yang, Jun Sun, Wei Li, Hongbo Yang, Linjie Wang, Fengying Gong, Shi Chen, Lin Lu, Hui Miao, Xianxian Yuan, Hanting Liang, Ran Li, Hui Huang, Zhiyu Peng, Asan, Hui Pan

## Abstract

Short stature is among the most common reasons for children being referred to the pediatric endocrinology clinics. The cause of short stature is broad, in which genetic factors play a substantial role, especially in primary growth disorders. However, identifying the molecular causes for short stature remains as a challenge because of the high heterogeneity of the phenotypes. Here, whole exome sequencing (WES) was used to identify the genetic causes of short stature with unknown etiology for 20 patients aged from 1 to 16 years old. The genetic causes of short stature were identified in 9 of the 20 patients, corresponding to a molecular diagnostic rate of 45%. Notably, in 2 of the 9 patients identified with genetic causes, the diagnosed diseases based on WES are different from the original clinical diagnosis. Our results highlight the clinical utility of WES in the diagnosis of rare, high heterogeneity disorders.

## INTRODUCTION

Short stature is one of the common chief complaints referred for genetics consultation in China. Medically, short stature is defined as a height-for-age less than two standard deviations (SD) below the average for a specific gender and race [1]. The cause of short stature is broad, which includes pathologic and non-pathologic factors. Short stature is classified into primary growth disorders, secondary disorders, and idiopathic short stature (ISS) by international agreement. Patients with ISS have no evidence of systemic, endocrine, nutritional, or chromosomal abnormalities, and have normal birth weight and sufficient growth hormone. Genetic factors play a pivotal role in short stature, especially in primary growth disorders. The genetic causes for short stature can be associated with chromosomal abnormalities, monogenic mutations, or complicate syndromes [1, 2].

Molecular genetic testing for short stature can help clinicians to provide specific diagnosis and prognosis and give recurrence risk assessment for patients and their family members. However, traditional genetic testing method for short stature based on Sanger sequencing technology is highly labor-intensive, costly, and time-consuming because of a limited number of genes tested each time. The success of traditional genetic testing to identify the disease-causing mutations highly depends on the accurate diagnosis in pre-test clinical evaluation, which determines the candidate gene list to be tested.

Next generation sequencing (NGS) technologies are high-throughput sequencing analysis, producing large-scale sequence data in one time. Targeted gene panel and whole exome sequencing based on NGS are two prevalent methods in clinical practice to identify the genetic etiology of inherited diseases, especially for the disorders with high genetic heterogeneity. WES, which simultaneously tests for nearly all coding regions in human genome (∼1∼2% of the genome, which harbors ∼85% of the known disease-causing mutations[3]), is an efficient strategy to determine the genetic basis of undiagnosed inherited diseases. It has been successfully used to diagnose undiagnosed cases of intellectual disabilities, other neurological diseases, developmental delay diseases, mitochondrial disease[4-8], and also short stature[9]. The general diagnostic rate ranged from 16% to 50%. Here, we analyzed 20 undiagnosed cases with short stature by WES and subsequently identified the genetic cause for short stature in 9 patients, which highlights the clinical utility of WES in the diagnosis of the highly heterogeneous disorders.

### Materials and methods

#### Patients

This study was approved by the Ethics Committee of Peking Union Medical College Hospital. All participants signed written informed consents. To determine the genetic causes, a cohort of subjects which was unable to identify the etiology of their short stature after receiving prior standard clinical diagnostic procedure was recruited from 2014 to 2015. Twenty patients with moderate short stature (2SD-3SD below the mean for age and sex) were selected for WES (patient’s records see Table 1). However, individuals were allowed to have additional phenotypes involving multiple systems or other hormonal deficiencies. Almost all patients are sporadic cases without family history except for patient 17 and patient 6. Case 17 has an affected sister with similar phenotype and mother of case 6 has bone abnormalities.

**Table 1.**
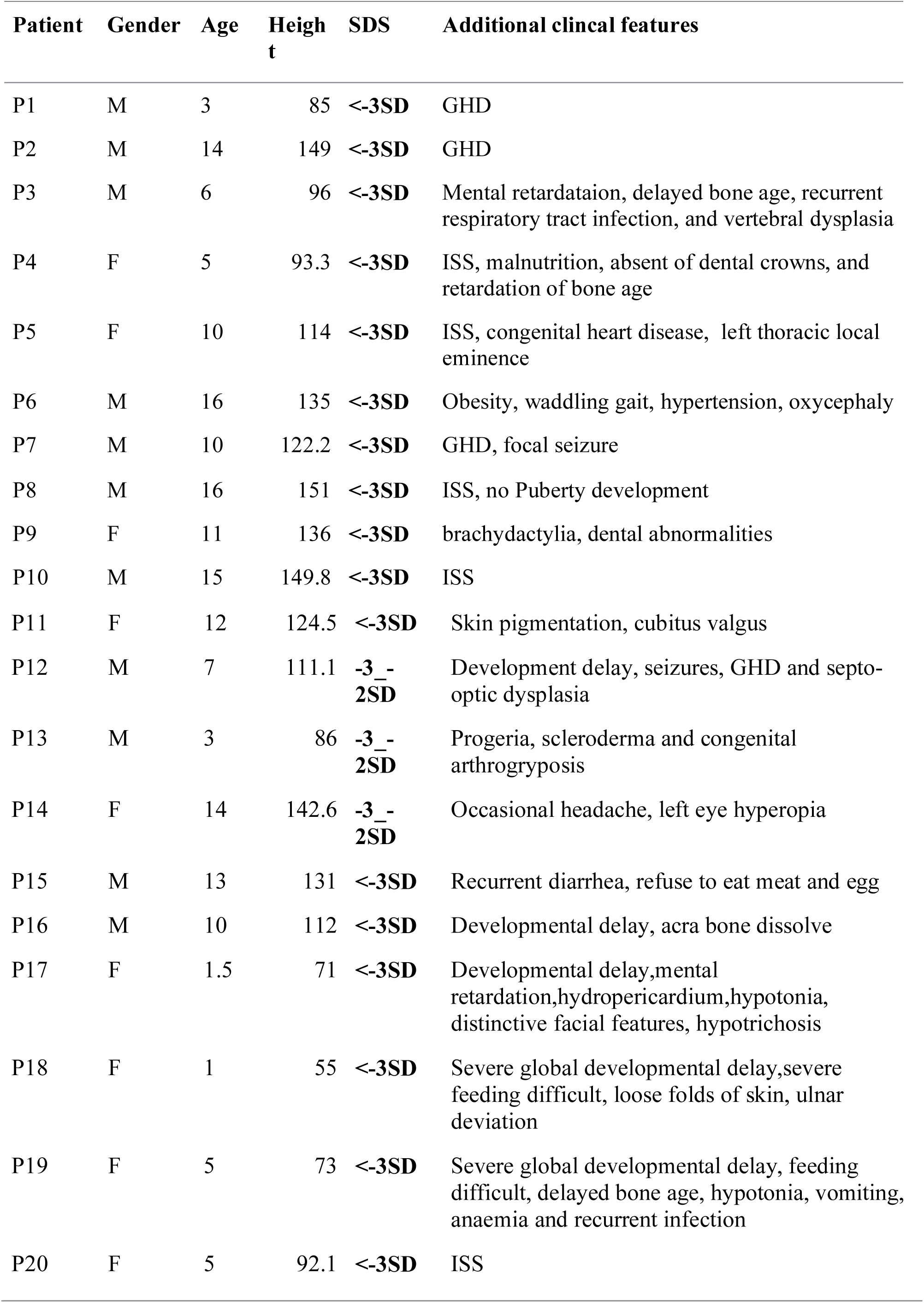
Patient clinical information overview

### Whole Exome Sequencing

Whole Exome Sequencing was performed as reported previously[10]. The variants interpretation was based on the workflow in Figure 1. The software used for non-synonymous functional predictions in the annotation pipeline include PolyPhen-2[11], SIFT[12] and Ens Condel[13]. Each variant was evaluated and categorized according to ACMG guidelines [14]. A primary gene list related with short stature including 469 genes was generated by using key word of short stature as a search term both in “disease features” and “text of disorders” in the Online Mendelian Inheritance in Man (OMIM) database (Supplementary Table 1). Sanger sequencing of all reported variants was performed for the proband as well as the parental samples, if available. Primers for PCR amplification were designed (Supplementary Table 2).

**Figure 1.**
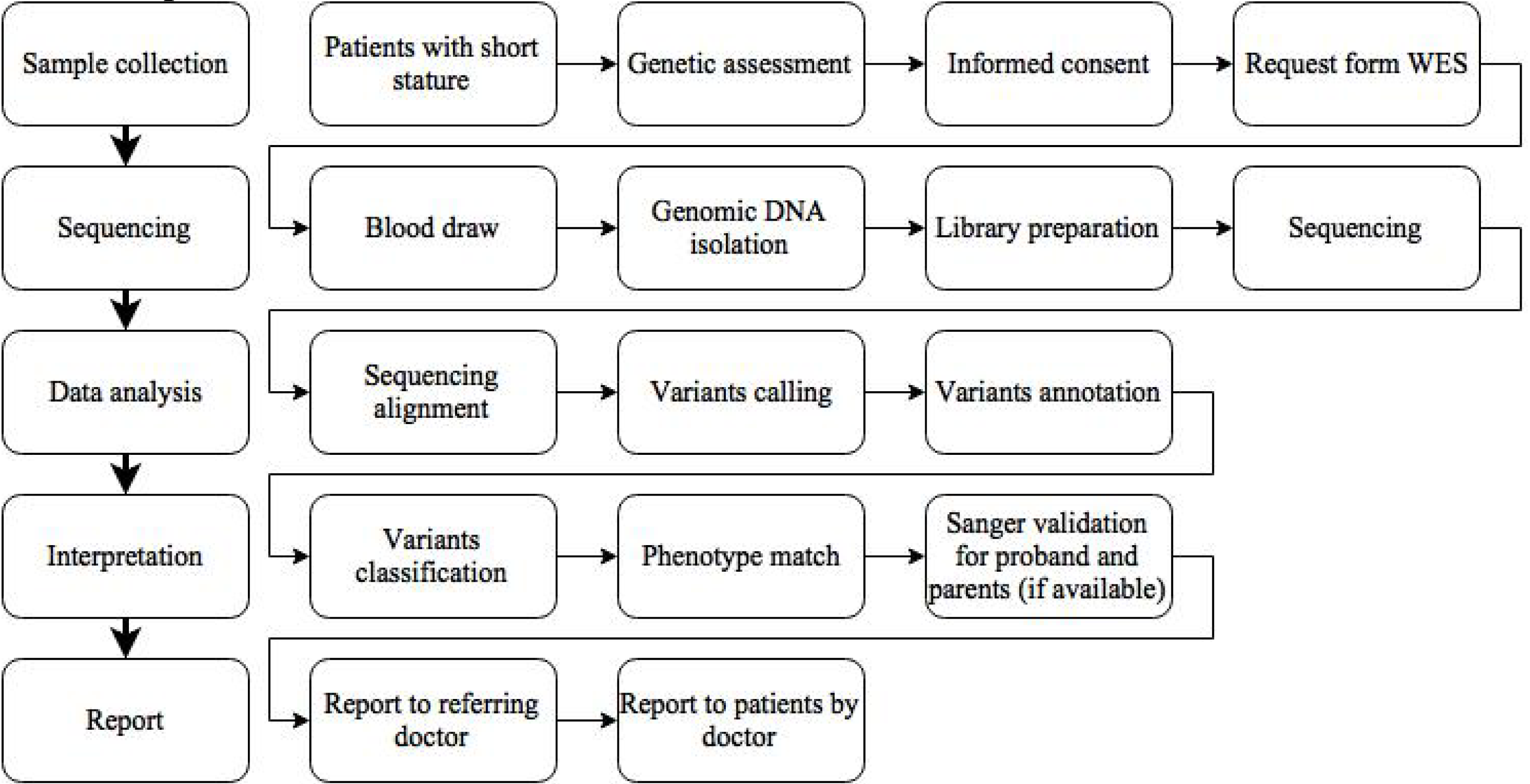
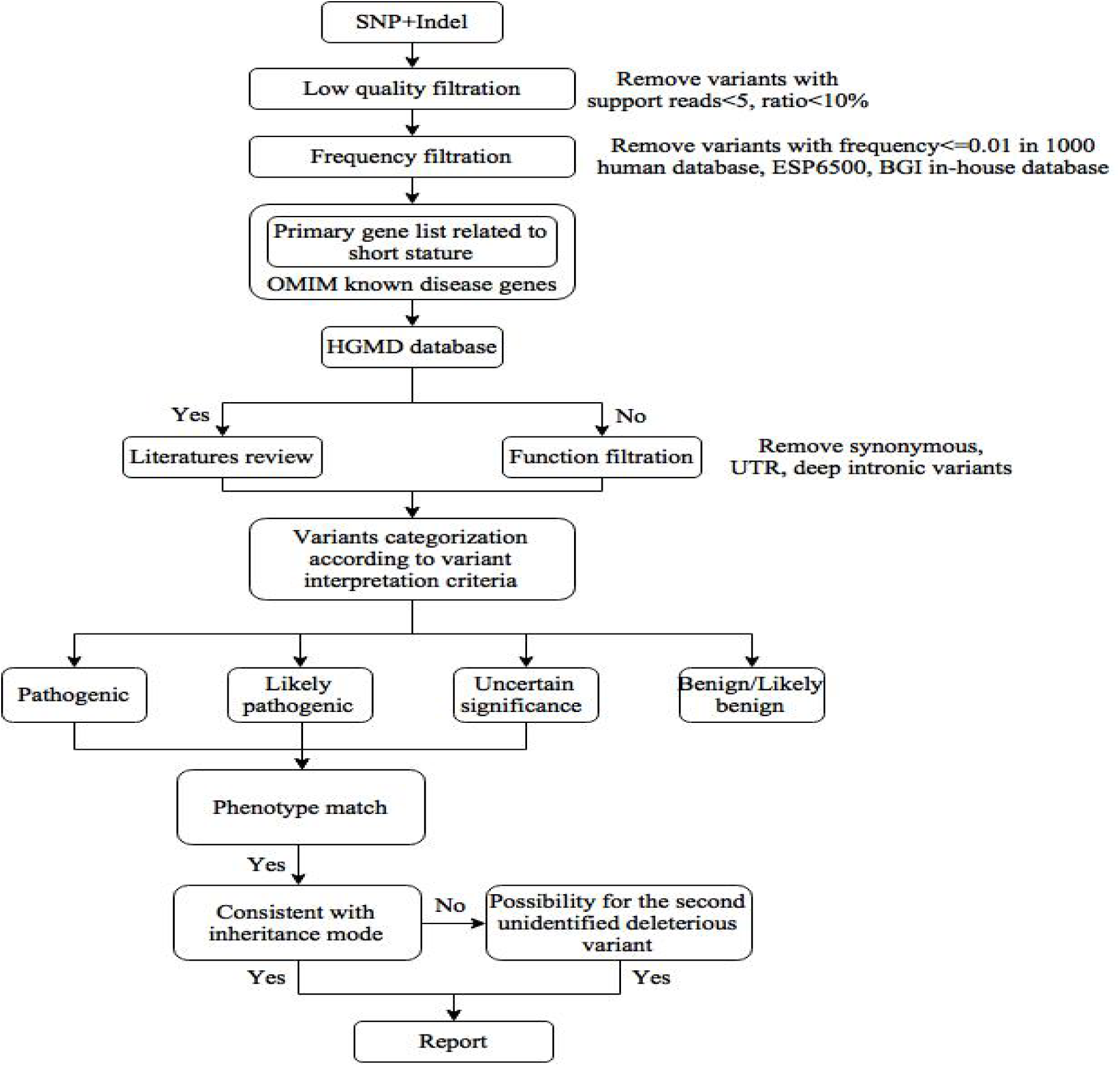

## Result

### Characteristics of the patients

All 20 patients are children or adolescents under 16 years, in which 11 are male and 9 are female. The ethnic of all patients are Chines Han. The heights of 17 patients are less than 3 standard deviations (SD), and the heights of 3 patients are between 2 to 3 SD below the mean height of corresponding age and sex group in the population. All the cases have received routine clinical diagnostic tests and are suspicious to have monogenic mutation genotype. Four patients have unexplained growth hormone deficiency (GHD), in which 1 patient is accompanied with septo-optic dysplasia (SOD). Six patients are diagnosed as idiopathic short stature (ISS), in which 1 patient is accompanied with congenital deafness. The remaining 10 patients are syndromic short stature with other abnormalities involving multiple systems such as skeletal deformity, skin abnormalities, seizure and mental retardation.

### Exome sequencing and potential pathogenic mutations

The data output and coverage statistics for each sample are shown in Supplementary Table 3. Generally, 10 GB raw data were generated for each patient, providing an average of 75 million sequencing reads with a 100-bp length mapped to target region and a 121-fold mean depth across the target region. Also, the average coverage at >10× read depth of the exome was 97%, which is within the expected coverage and depth for WES studies. An average of 20,536 single-nucleotide variants were identified in each exomes. After function and frequency filtration, about 145 variants of potential clinical significance on known disease-causing genes recorded in OMIM database were retained to be interpreted for each sample, of which about 5 variants were recorded in Human Gene Mutation Database (HGMD) (Table 2), and 20 variants were from genes associated with short stature.

**Table 2.**
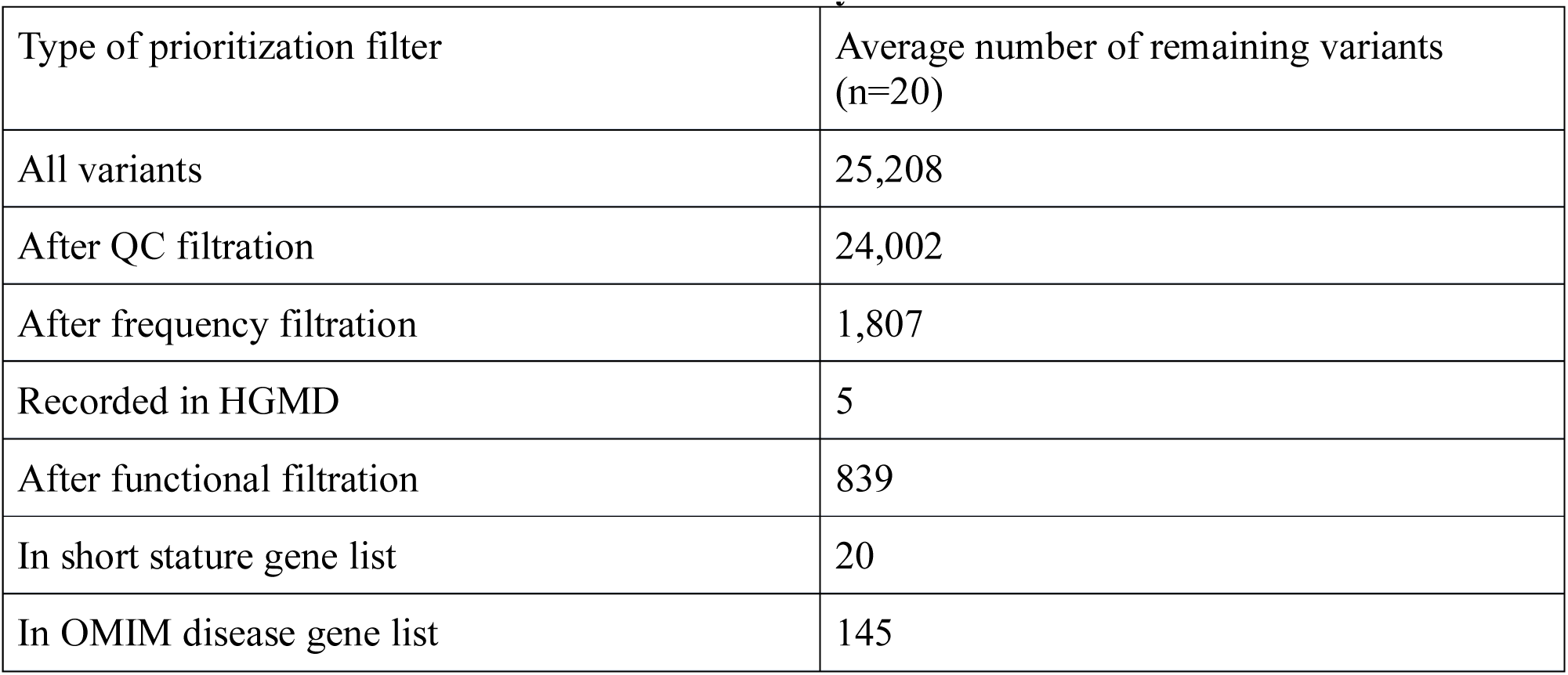
Prioritization scheme for exome data analysis

Of 20 probands, 2 homozygous mutations and 7 heterozygous mutations were identified in 9 patients under the criteria for a molecular diagnosis. The positive molecular diagnostic rate in this 20 patients was 45%. All positive cases met all of the diagnostic criteria corresponding to their mutation types, appropriate inheritance patterns (if data available), and disease-phenotype relationship. Most of the identified mutations are known to be pathogenic except 2 novel mutations. In 9 identified mutations, 5 are missense mutations, 2 are nonsense mutations, 1 is frameshift mutation, and 1 is synonymous mutation (Table 3). To validate the variants identified by WES, Sanger sequencing was performed for the proband as well as the parental samples, if available (Supplementary Figure 1).

**Table 3.** Summary of identified potential pathogenic mutation

| Gene | Transcript ID | Neucleotide change | Amino Acid change | Zygosity | function | rs ID | PolyPhen2 | Condel | Categori |
| --- | --- | --- | --- | --- | --- | --- | --- | --- | --- |
| COL2A1 | NM_001844.4 | c.1124G>T | p.Gly375Val | Het | Missense | -l | - | - | Likely Pathoger |
| ACAN | NM_013227.3 | c.6240_6244del TGGAG | p.Gly2081Tyrfs*18 | Het | Frameshift | - | - | - | Likely Pathoger |
| LMNA | NM_170707.3 | c.1824C>T | p.Gly608Gly | Het | Synonymous | rs58596362 | - | - | Pathoger |
| SLC7A7 | NM_001126106.2 | c.725G>A | p.Trp242* | Hom | nonsense | - | - | - | Likely Pathoger |
| PRKAR1A | NM_002734.3 | c.1102C>T | p.Arg368* | Het | nonsense | rs387906692 | - | - | Pathoger |
| PTPN11 | NM_002834.3 | c.922A>G | p.Asn308Asp | Het | Missense | rs28933386 known | 0.329 | Neutral | Pathoger |
| LMNA | NM_170707.3 | c.1580G>A | p.Arg527His | Hom | missense | rs57520892 | 0.051 | Neutral | Pathoger |
| BRAF | NM_004333.4 | c.770A>G | p.Gln257Arg | Het | missense | rs180177035 | 0.456 | Deleterious | Pathoger |
| HRAS | NM_001130442.1 | c.34G>T | p.Gly12Cys | Het | missense | rs104894229 | 0.899 | Deleterious | Pathoger |

### Diagnosis based on whole-exome sequencing

#### Patient 6

In addition to short stature, the clinical features of Patient 6 include osteoporosis of spine and pelvis, multiple skeletal deformities, and arthrogryposis. This patient, whose mother has similar skeletal abnormalities, was clinically diagnosed to be Sondyloepiphyseal Dysplasia Congenital (SEDC, OMIM 183900). A heterozygous novel possibly pathogenic mutation c.1124G>T was detected on the *COL2A1* gene. *COL2A1* gene encodes a 134.4 kDa protein called α-1 chain of collagen type II with 1,487 amino acids. The three identical polypeptide chains (Gly-X-Y) is the backbone of the Type II collagen. Most patients with SEDC carry either glycine substitutions destroying the Gly-X-Y triplets or splice site mutations causing in-frame deletions. c.1124G>T is a missense mutation from a helical Gly to Val substitution at codon 375 in a Gly-X-Y repeat. Moreover, at the same codon, Gly to Asp substitution of *COL2A1* gene was identified in a case with SEDC at adult age [14], so c.1124G>T is likely to disrupt the correct folding of Type II collagen. The mutation c.1124G>T is not detected in any sequenced databases (i.e. samples in the exome sequencing project, dbSNP, 1000 Genomes databases, and BGI in-house database). Mutation of *COL2A1* gene may cause a spectrum of skeletal disorders. In OMIM database, there are 15 phenotypes related with *COL2A1* gene, one of which is SEDC. The disorder is characterized by disproportionate short-trunk dwarfism, skeletal and vertebral deformities, including scoliosis and thoracic hyperkyphosis, coxa vara, avascular necrosis-like changes in bilateral femoral epiphyses, hip pain or waddling gait, genu valgum, and various joint diseases[15, 16]. The clinical features of SEDC well matched with the phenotype and original diagnosis of Patient 6. A heterozygous mutation c.1124G>T of *COL2A1* gene was identified in his mother who has the similar skeletal abnormalities by Sanger sequencing (Supplementary Figure 1) and this is consistent with the autosomal dominant inheritance pattern of SEDC caused by mutation of *COL2A1* gene. c.1124G>T of *COL2A1* gene was assumed to be the disease-causing mutation of this patient.

#### Patient 9

Patient 9 is an 11-year-old girl born SGA (full-term birth weight 2.3kg). The patient presented to have short stature, lower limb shortening, brachydactyly and dental abnormalities. A heterozygous null mutation c.1102C>T of *PRKAR1A* gene was detected in this patient. This mutation causes amino acid at position 368 to change from Arg to stop codon, which is predicted to result in premature termination. Moreover, several cases with mutation c.1102C>T of *PRKAR1A* gene have been reported. Linglar et al. [17]identified a *de novo* heterozygous c.1102C>T transition in exon 11 of the *PRKAR1A* gene in 3 unrelated patients with acrodysotosis-1 with hormone resistance (ACRDYS1; OMIM 101800). In 4 unrelated patients with acrodysostosis with hormone resistance, Michot et al. [18] identified a heterozygous de novo R368X mutation. All of these patients had short stature, severe brachydactyly, and other bone abnormalities, which are pretty similar to patient 9.

#### Patient 10

Patient 10 was diagnosed as ISS without any obvious abnormality. A heterozygous novel frameshift mutation c.6240_6244delTGGAG of *ACAN* gene was identified in this patient. Deletion of 5 nucleotides in exon 12 changes the reading frame of *ACAN* mRNA and leads to a premature termination downstream of codon 2081. This mutation was not found in the sequenced population genomes or exomes. In OMIM database, ACAN gene is known to cause three diseases including Osteochondritis dissecans, short stature and early-onset osteoarthritis (Phenotype MIM number 165800), Spondyloepimetaphyseal dysplasia, aggrecan type (Phenotype MIM number 612813), and Spondyloepiphyseal dysplasia, Kimberley type (Phenotype MIM number 608361). The phenotype presented in all three diseases includes various skeletal abnormalities which are not observed in this patient. However, loss of function in the aggrecan (*ACAN*) gene has been identified by WES in three families with short stature and advanced bone age recently. The study draws a conclusion that mutations in the *ACAN* gene should be included in the differential diagnosis of children with idiopathic short stature or familial short stature[19]. After reviewing the supporting evidence above mentioned, we assumed the mutation c.6240_6244delTGGAG of *ACAN* gene as the cause of ISS in this patient.

#### Patient 11

Patient 11 is a 12-year-old girl who was full-term born without any obviously abnormalities. The phenotype of this patient includes short stature, retardation of bone age, developmental delay, skin pigmentation, and cubitus valgus,without typical facial characteristics. A heterozygous mutation c.922A>G in PTPN11 gene was identified in this patient. c.922A>G was the most common disease causing mutation for Noonan syndrome (NS), and also a hotspot mutation of *PTPN11* [20-22]. NS is a clinically heterogeneous disorder, which is featured with short stature, congenital heart defect, and developmental delay of various degrees. Other features may include broad or webbed neck, unusual chest shape with superior pectus carinatum and inferior pectus excavatum, cryptorchidism, characteristic facies, varied coagulation defects, lymphatic dysplasias, and ocular abnormalities[23]. We have noticed that Patients 11, as well as other patients carrying mutation c.922A>G (Asn308Asp), received normal education.

#### Patient 13

This patient was clinically diagnosed to be progeria. Clinical features of this patient include short stature, scleroderma, and congenital arthrogryposis. A known pathogenic mutation c.1824C>T of *LMNA* gene was detected. *LMNA* is known to cause the Hutchinson-Gilford progeria syndrome (HGPS), which is a rare genetic disorder featured with many symptoms of premature ageing. There is a considerable overlap in clinical features between HGPS and Patient 13. Most cases of HGPS refer to a heterozygous synonymous mutation (c.1824C>T; p.Gly608Gly) which increases the possibility using an internal 5’ splice site (5’SS) in exon 11 of the *LMNA* pre-mRNA and results in a truncated protein (progerin) with a dominant negative effect [24]. So c.1824C>T is assumed to be the cause of the disease in this patient.

#### Patient 18

Patient 18 is an one-year-old girl born with normal weight and length. The patient presented with severe delayed development, hypotonia, various skeletal abnormalities, loose skin, sparse hair, and extremely difficulty feeding. The baby didn’t gain any weight during the year after birth. A de novo heterozygous mutation c.34G>T of *HRAS* gene was detected in this patient. c.34G>T of *HRAS* was a known pathogenic mutation which can cause Costello Syndrome (OMIM 218040). Nucleotide 34 encoding the glycines at position 12 is the mutation hotspot of *HRAS* gene. Most of the reported Costello Syndrome patients were caused by this mutation [25-28]. Costello syndrome is a rare congenital condition associated with short stature, characteristic coarse faces, distinctive hand posture and appearance, severe feeding difficulty and growth retardation in all cases [25]. The phenotypes of this patient matched well with that of reported Costello syndrome patients. c.34G>T is de novo because this mutation was not detected in other unaffected family members (parents and brother) by Sanger sequencing (Supplementary Figure 1), which was consistent with the dominant inheritance pattern of Costello syndrome caused by the mutation of *HRAS* gene.

#### Patient 17

Besides of short stature, the additional clinical features of Patient 17 include severe developmental delay, mental retardation, hydropericardium, hypotonia, distinctive facial features, and hypotrichosis. A de novo heterozygous missense mutation c.770A>G of *BRAF* gene was detected in this patient. c.770A>G of *BRAF* causing a p.Gln257Arg amino acid change has been reported previously in literatures as the cause of cardiofaciocutaneous syndrome 1 (CFC1, OMIM 115150) [18]. Many body systems were involved in patients of cardiofaciocutaneous syndrome, particularly the heart (cardio-), facial features (facio-), and the skin and hair (cutaneous). The major features of CFC include characteristic craniofacial dysmorphology, congenital heart disease, dermatologic abnormalities, failure to thrive, and amentia [18]. The disease features of cardiofaciocutaneous syndrome can be overlapped with most of the clinical phenotype of this patient. Sanger validation of c.770A>G of *BRAF* gene for the parental samples showed that both unaffected parents were negative for this mutation, which was consistent with the dominant inheritance mode of CFC1.

### Corrected patient diagnoses

#### Patient 16

This patient presented acroosteolysis, colored skin, and short stature, and was diagnosed to have Hajdu-Chany Syndrome. Sanger sequencing for all the exons of *NOTCH2* gene related with Hajdu-Chany Syndrome was performed and no pathogenic mutation was identified in this patient. A homozygous missense mutation c.1580G>A of *LMNA* gene was identified. c.1580G>A is a known pathogenic mutation which can cause Mandibuloacral Dysplasia (MADA, OMIM 248370). Affected individuals are undistinguishable at birth and then progressively present lipodystrophy, dysmorphic craniofacial, and skeletal features. Other features of MADA include mandibular hypoplasia, acro-osteolysis, prominent appearance of the eyes, dental overcrowding, beaked nose, delayed closure of the cranial sutures, clavicular dysplasia/osteolysis, joint contractures, and poikiloderma. This mutation has been detected in at least 10 unrelated MADA patients with similar clinical features [29-31]. Sanger validation of c.1580G>A was performed on the samples of family members including affected sister and unaffected parents, and the result showed that both parents are carrier of heterozygous c.1580G>A and the affected sister is homozygous c.1580G>A which is identical with the patient. This result is consistent with the autosomal recessive inheritance pattern of MADA. We concluded that the homozygous mutation c.1580G>A of *LMNA* gene is possibly the cause of disease in this patient. The original diagnosis for this patient was Hajdu-Chany Syndrome. Whole-exome sequencing test successfully corrected the diagnosis.

#### Patient 15

Patient 15 was a 13-year-boy who was full-term born by normal spontaneous vaginal delivery. He presented developmental delay since 4 months old, and refused to eat egg and meat. The patient has poor appetite since infancy and suffers from perennial diarrhea. A homozygous null mutation c.725G>A of *SLC7A7* was identified. Although c.725G>A is a novel mutation which has not been reported previously, it is a severe loss of function mutation which results in a premature termination at codon 242. A mutation in the same codon c.726G>A which also results in a premature termination at codon 242 has been reported [29] to be the disease-causing mutation in 2 unrelated patients with lysinuric protein intolerance (OMIM: 222700). The clinical features are similar between this patient and other reported patients with this disease. The original diagnosis for this patient was ISS. The WES result showed that this patient was affected by lysinuric protein intolerance, which meant that the short stature and developmental delay was one of the complications caused by malnutrition.

## Discussion

Short stature is among the most common reasons for children visiting pediatric endocrinology clinics in China. Some cases are non-familial short stature without hormonal deficiency and other identified causes and diagnosed as idiopathic short stature (ISS). Molecular genetic testing can help the clinicians to provide specific diagnosis, prognosis, and give recurrence risk assessment for patients and their family members. Furthermore, the genetic diagnosis can help clinicians to efficiently manage and treat many of these patients. However, high genetic and clinical heterogeneity of short stature challenges the traditional Sanger sequencing test. WES has the advantage of speed and efficiency in identifying the disease-causing mutation of short stature by sequencing the coding regions of whole genome simultaneously using next-generation sequencing techniques.

We obtained molecular diagnosis for 9 patients through applying WES on the 20 unselected short stature patients (Table 4). The results of molecular diagnosis for Patient 15 and Patient 16 were different from the initial diagnosis, which indicated that WES can not only confirm the diagnosis, but also correct the clinical diagnosis. The total diagnose rate of this study is 45%. Although the scale of the study population are not big enough to yield a diagnostic rate with statistical significance, it is still a strong evidence highlighting the clinical utility of WES for patients with unexplained short stature, and also for other types of diseases with high heterogeneity like short stature.

**Table 4.**
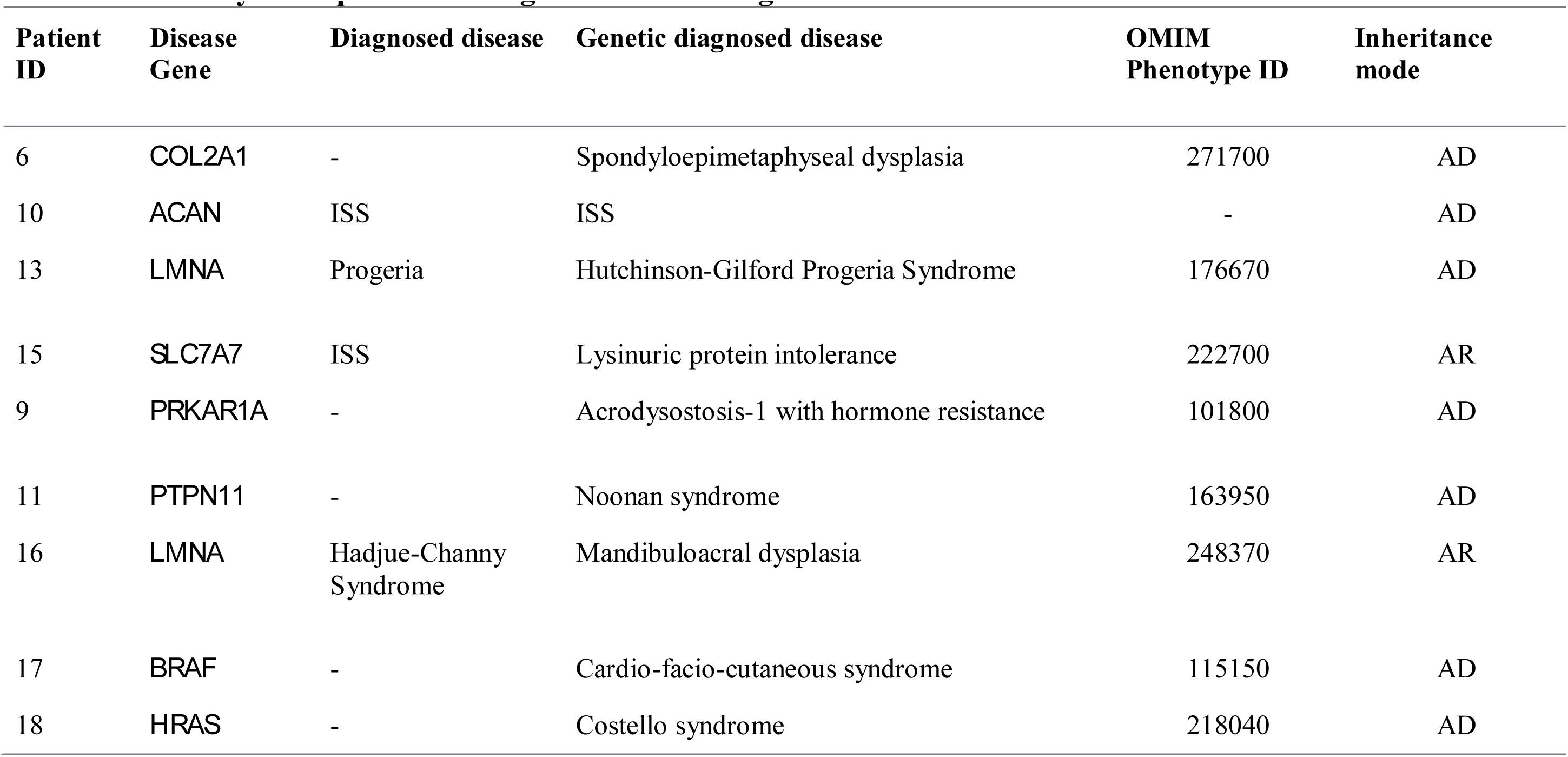
Summary of 10 patients who get molecular diagnosis

For the past few years, several clinical applications of WES on genetic disease have been reported. The reported success rates vary between 25% and 30% [4, 6], which is lower than the 45% we got. A possible reason is that the patients in our study are all directly referred from the clinicians without any genetic test. A paper has been published by ACMG [30] to provide a rubric for the evaluation of short stature. In the guideline, genetic tests are recommended to be performed firstly for short stature patients. For the short stature patients with other recognized syndromes, there should be specific testing that includes related genes. In some WES studies, some patients with specific or recognizable genetic cause have already accepted some genetic tests to exclude the most possible reason, which will decrease the diagnosed ratio of WES test.

Although WES is an ideal technology for the disorders with high heterogeneity like short stature, it is still a challenge to develop an effective interpretation workflow to select the real disease-causing mutations from about 25,000 variants. To limit the number of variants needed to be reviewed, a priority interpretation workflow was selected to identify potential candidate variants. The function filtration can help to remove the variants with no effects on protein. The non-effective variants usually include silent variants and variants that occur in deep intron or untranslated region (UTR), and some of which have also been reported as pathogenic mutations because of the disruption or introduction of splice site. In Patient 13, a known pathogenic synonymous mutation on *LMNA* gene was detected to be the cause of disease, which suggested that it is necessary to perform the known database search before the function filtration removes non-effective variants.

There are several reasons for the failure to find the molecular etiology in the patients. The first one is that the patients are affected with disease of currently unknown genetic causes. Although WES can capture the exome region within the whole genome, our interpretation is based on the known disease-causing genes. We will further analyze the data to identify new disease-causing genes, but we also have to consider the possibility of the disease not being mendelian. The second reason is the limitations of WES technology. Large duplication and deletion, balanced translocation, inversions, SNP in the regulation and deep intron, ploidy changes, uniparental disomy, and methylation alterations cannot be detected by WES. It is likely for the pathogenic mutations to occur in these regions without being detected by WES.

“Incidental” or “off-target” findings remain an area of debate in medical genetics. Our study focused on the genetic diagnosis of related diseases, so incidental findings are not included in the report sent to patients or their guardians.

In conclusion, a workflow based on WES that identifies the genetic cause of short stature has been developed in our study. By applying WES to the 20 unselected short stature patients with no confirmed diagnosis, we got a 45% diagnose rate. The results proved our approach effective for identifying the genetic cause of short stature, and also highlighted the clinical utility of WES in short stature and other heterogeneity diseases.

For the patients who failed to be identified with any genetic cause, the data will be reanalyzed based on the updated known disease-causing genes list after a period of time. We will aim to determine new disease-causing genes in short stature patients who still remain undiagnosed.

## Acknowledgement

This work was funded by the National Key Research and Development Program of China (2016YFC0901501),and CAMS Innovation Fund for Medical Science(CAMS-2016-12M-002). We thank all patients and their families for participating in the project. We thank our colleagues from the genome sequencing platform in BGI-Tianjin for assistance in sequencing.

